# Combinatorial mRNA vaccination enhances protection against SARS-CoV-2 delta variant

**DOI:** 10.1101/2021.12.08.471664

**Authors:** Renee L. Hajnik, Jessica A. Plante, Yuejin Liang, Mohamad-Gabriel Alameh, Jinyi Tang, Chaojie Zhong, Awadalkareem Adam, Dionna Scharton, Grace H. Rafael, Yang Liu, Nicholas C. Hazell, Jiaren Sun, Lynn Soong, Pei-Yong Shi, Tian Wang, Jie Sun, Drew Weissman, Scott C. Weaver, Kenneth S. Plante, Haitao Hu

**Author notes:** Correspondence Haitao Hu, PhD, Kenneth S. Plante, PhD. These authors contribute equally (Renee L. Hajnik, Jessica A. Plante, Yuejin Liang).

## Abstract

Emergence of SARS-CoV-2 variants of concern (VOC), including the highly transmissible delta strain, has posed challenges to current COVID-19 vaccines that principally target the viral spike protein (S). Here, we report a nucleoside-modified mRNA vaccine that expresses the more conserved viral nucleoprotein (mRNA-N). We show that mRNA-N alone was able to induce a modest but significant control of SARS-CoV-2 in mice and hamsters. Critically, by combining mRNA-N with the clinically approved S-expressing mRNA vaccine (mRNA-S-2P), we found that combinatorial mRNA vaccination (mRNA-S+N) led to markedly enhanced protection against the SARS-CoV-2 delta variant compared to mRNA-S. In a hamster model, we demonstrated that while mRNA-S alone elicited significant control of the delta strain in the lungs (∼45-fold reduction in viral loads compared to un-vaccinated control), its effectiveness in the upper respiratory tract was weak, whereas combinatorial mRNA-S+N vaccination induced markedly more robust control of the delta variant infection in the lungs (∼450-fold reduction) as well as in the upper respiratory tract (∼20-fold reduction). Immune analyses indicated that induction of N-specific immunity as well as augmented S-specific T-cell response and neutralizing antibody activity were collectively associated the enhanced protection against SARS-CoV-2 delta strain by combinatorial mRNA vaccination. These findings suggest that the combined effects of protection in the lungs and upper respiratory tract could both reduce the risk of severe disease as well as of infection and transmission.

## INTRODUCTION

COVID-19, caused by SARS-CoV-2, has spread rapidly and led to a major global pandemic since its detection in December 2019 [1, 2]. A number of vaccines based on various platforms have been developed in response to the COVID-19 pandemic, including DNA [3, 4], mRNA [5-8], viral vectors [9-17], protein subunit [18-20], and inactivated vaccines [21], among which the two mRNA vaccines and the Ad26-vectored vaccine showed high efficacy in late-stage clinical trials and have thus been licensed or received emergency use authorization (EUA) in the US and many other regions of the world [22, 23]. While rapid development of these vaccines provides hope for ending the COVID-19 pandemic, emergence of variants of concern (VOCs), including the highly transmissible delta strain, has posed constant challenges for vaccine-induced immunity [24-28]. For example, multiple spike variants have been identified that show reduced sensitivity to neutralization by vaccine-induced humoral immunity [26-28]. Clinical studies also indicate that the two approved mRNA-S vaccines (BNT162b2 and mRNA-1273), while retaining high effectiveness in protecting against diseases caused by the delta variant, show lower overall efficacy in preventing infection [29].

Current COVID-19 vaccines principally target the viral spike protein (S), or its receptor-binding domain (RBD) in particular, with the major goal of eliciting a potent neutralizing antibody response [22, 23]. We hypothesize that vaccine approaches targeting a relatively more conserved viral protein in addition to the S protein would likely provide broader protection, especially against VOCs. Among the SARS-CoV-2 proteins, the nucleoprotein (N) is another dominant antigen to trigger host immune response [30, 31] and is more conserved across different SARS-CoV-2 variants as well as different coronaviruses compared to the S protein [32]. Prior evidence has suggested that besides the S protein, N can induce long-lasting and broadly reactive T cells [33] that correlate with control of coronavirus [31]. Thus, the N protein may represent a promising immunogen for incorporation in the SARS-CoV-2 vaccine design [16, 34] to provide a more robust vaccine platform in face of future viral mutations.

In this study, we generated a nucleoside-modified (m1Ψ) mRNA vaccine that encodes the SARS-CoV-2 N protein (mRNA-N) and is formulated in lipid nanoparticles (LNP). We showed that mRNA-N was highly immunogenic and induced robust N-specific T-cell responses and binding IgG. As expected, no neutralizing antibody was elicited by the mRNA-N vaccine alone. In mice and hamsters challenged with SARS-CoV-2, we demonstrated that mRNA-N alone induced only modest but significant control of mouse-adapted SARS-CoV-2 strain as well as the delta strain (B.1.617.2) infection. Further, in the hamster model, we compared the protective efficacy of the combinatorial mRNA-S+N vaccination with the clinically approved S-expressing mRNA vaccine (mRNA-S-2P) alone on immune control of SARS-CoV-2 delta strain in both lungs and upper respiratory tract. While the mRNA-S alone led to ∼45-fold reduction in viral loads in the lungs compared to the mock-vaccinated controls, its effect against the delta strain in the upper respiratory tract was weak. Serum analyses indicated that neutralizing activity elicited by mRNA-S against delta strain was markedly reduced (by ∼5-fold in mean PRNT_50_) compared to that against WT virus. Notably, combinatorial mRNA-S+N vaccination not only induced robust N-specific T-cell immunity, but also augmented S-specific CD8 T-cell response and neutralizing antibody activity, leading to more robust control of the delta infection in the lungs (∼450-fold reduction) as well as in the upper respiratory tract (∼20-fold reduction). Thus, our study presents a new mRNA vaccine platform that provides stronger protection against the SARS-CoV-2 delta variant. The findings also indicate potential broad utility of this vaccine platform against all future VOCs.

## RESULTS

### mRNA-N vaccine generation and immunogenicity analysis

Since coronavirus N protein represents an important viral antigen to induce durable and broadly reactive T cells, we designed and generated a methyl-psuedouridine-modified (m1Ψ) mRNA that encodes the full-length N protein of SARS-CoV-2 (Wuhan-Hu-1 strain) (**Fig. S1a**). Synthesis, purification, and lipid nanoparticle (LNP) formation of the mRNA-N vaccine were conducted as previously described [35-37]. Expression of N protein in cells following mRNA-N transfection was confirmed by western blot (**Fig. S1b)**.

Immunogenicity of mRNA-N was evaluated in wild-type (WT) Balb/c mice. Two groups of mice (8/group) were vaccinated with PBS (mock) or mRNA-N (1µg). mRNA dose was selected based on previous studies in mice [38, 39]. Vaccination was given intramuscularly (i.m.) at week 0 (prime) and week 3 (booster) **(Fig. S2a)**. Three weeks after prime vaccination (on the day of booster), blood/sera were collected for analysis of antibody response; two weeks after booster vaccination (week 5), mice were euthanized and subjected to analyses of vaccine-induced humoral and cellular immune responses **(Fig. S2a)**. First, T-cell immunity was examined in splenocytes by flow cytometry. Based on CD44 expression, we observed that compared to the mock controls, mRNA-N vaccination induced activation of total CD4^+^ and CD8^+^ T cells in the spleen (**Fig. 1a**). Vaccine-induced N-specific T-cell response was examined by intracellular cytokine staining (ICS) and flow cytometry. Representative FACS plots for cytokine expression (IFN-γ, TNF-α, IL-2) in T cells following recall peptide stimulation are shown in **Fig. 1b** and **Fig. S2b**. Compared to the mock controls, mRNA-N vaccination induced high magnitudes of N-specific CD4^+^ and CD8^+^ T cell response in the spleen (p<0.0001 for all three cytokines) (**Fig. 1c-d)**. N-specific T cells appeared to predominantly express TNF-α (mean: 1.65% for CD4^+^ T cells and 0.83% for CD8^+^ T cells), followed by IFN-γ and IL-2 (**Fig. 1c-d)**. The mRNA-N vaccine induced T-cell response was further evaluated by IFN-γ ELISPOT (**Fig. S2c**) and it was confirmed that compared to the mock control, mRNA-N vaccine elicited high levels of N-specific T cells in spleen (mean SFC/10^6^ splenocytes for mock vs. mRNA-N: 8 vs. 637) (p<0.0001) (**Fig. 1e**).

**Figure 1.**
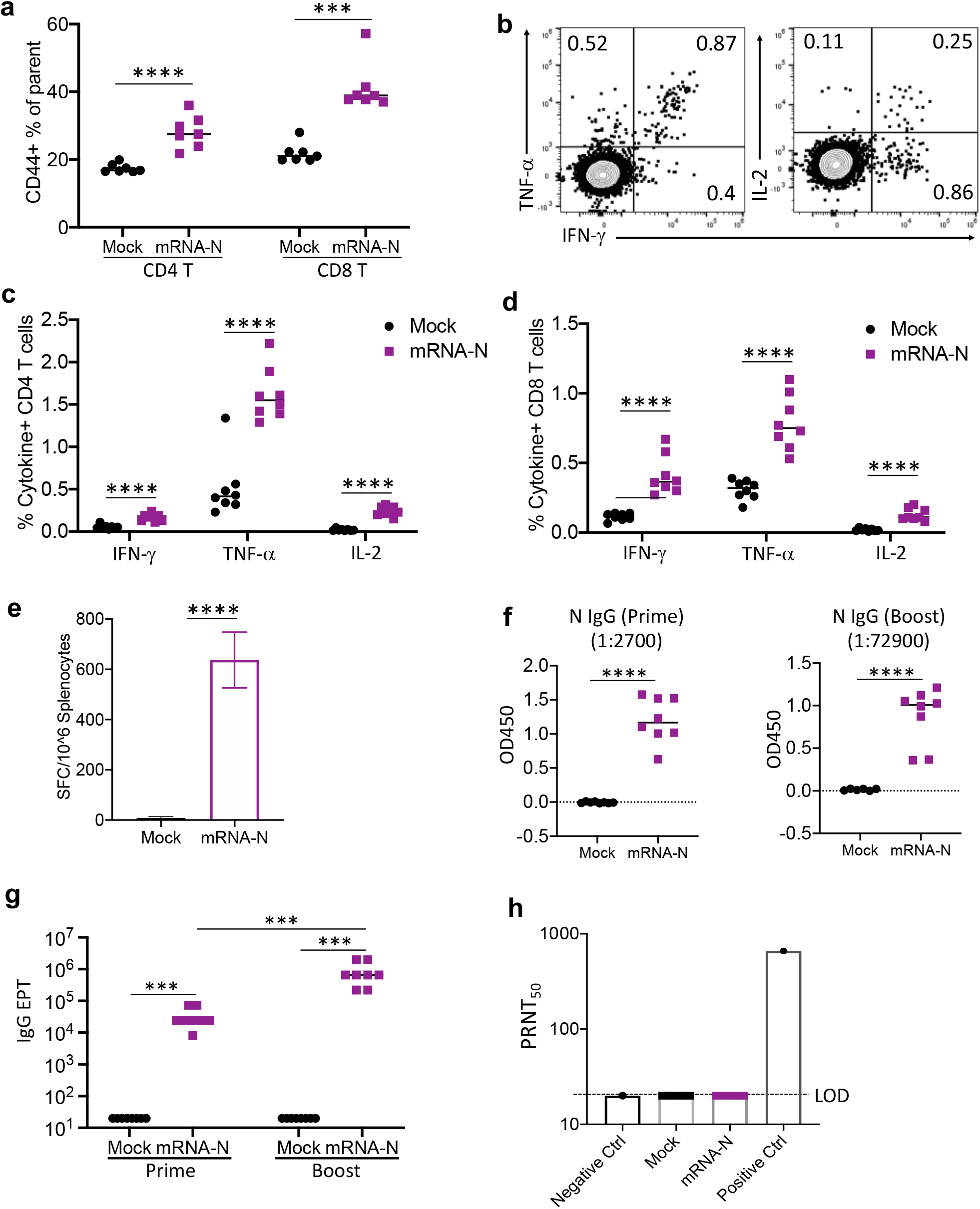
mRNA-N vaccine immunogenicity in mice. **(a)** Analysis of total CD4^+^ and CD8^+^ T cell activation in the mouse spleen following mock or mRNA-N vaccination. Splenocytes collected at week 5 (2 weeks after booster) were stained for mouse CD3, CD4, CD8, and CD44. Expression of CD44 on total CD4^+^ and CD8^+^ T cells were examined by flow cytometry and shown as % CD44^+^ in CD4^+^ or CD8^+^ T cells. **(b)** ICS measurement of vaccine-specific T cells in mouse spleen. Representative FACS plots for cytokine expression in T cells were shown. **(c-d)** Comparison of % cytokine-positive, N-specific CD4^+^ (c) or CD8^+^(d) T cells in the spleen between mock and mRNA-N vaccine groups. (**e**) Comparison of levels of N-specific T cells in the spleen measured by IFN-γ ELISPOT. Data were shown as spot forming cells (SFC) per 10^6^ splenocytes (mean ± SD). **(f)** ELISA measurement of serum N-specific binding IgG following prime (week 3) or booster (week 5) vaccination. OD values for individual serum samples after prime or booster vaccination at the indicated serum dilution (1:2700 for prime; 1:72900 for booster) were shown. **(g)** Comparison of N-specific binding IgG endpoint titers (EPT) between mock and vaccine groups after prime and booster vaccination. **(h)** Serum neutralizing activity measured by Plaque Reduction Neutralization Test (PRNT) using WT SARS-CoV-2. PRNT_50_ for individual serum samples of the mock and vaccine groups were shown. LOD: limit of detection. Negative control and positive control were included. One-way ANOVA or Student’s t-test were used for statistical analysis. * p<0.05, ** p<0.01, *** p<0.001, **** p<0.0001.

We next examined mRNA-N-induced antibody responses in the mouse sera following prime and booster immunization. First, ELISA was performed to examine N-specific binding IgG. Compared to mock control, prime immunization with the vaccine induced significant binding IgG (high OD values for sera diluted at 1:2700) (**Fig. 1f**; left panel), which was markedly enhanced by booster immunization (high OD values for sera diluted at 1:72900) (**Fig. 1f**; right panel). To determine antibody endpoint tiers (EPT), sera were serially diluted and N-specific binding IgG for each mouse sample was examined by ELISA (**Fig. S2d**). The analysis showed that median IgG EPTs after prime and booster vaccination were 24300 and 656100, respectively (**Fig. 1g**). Finally, serum neutralizing activity was determined by the Plaque Reduction Neutralization Test (PRNT). As expected, based on the lack of exposure to the S protein, no neutralizing activity was detected in any of the vaccinated animals **(Fig. 1h**). Together, these data suggest that the mRNA-N vaccine is highly immunogenic and i.m. immunization induces robust N-specific T-cell immunity and binding antibody response.

### mRNA-N vaccine alone induces modest but significant control of SARS-CoV-2 in mice and hamsters

Because it remained unclear if immunization with N-expressing vaccine alone would induce immune control of SARS-CoV-2, we evaluated the effectiveness of mRNA-N vaccine in animal models. First, two groups of WT Balb/c mice (8/group) were similarly vaccinated with either PBS (mock) or mRNA-N vaccine as described above at week 0 (prime) and week 3 (boost), followed by intranasal challenge with a mouse-adapted SARS-CoV-2 strain (MA-SARS-CoV-2; 2×10^4^ pfu) [40] at week 5 (**Fig. S3a**). Two days post-infection (2 DPI), viral RNA copies in the lung were measured by quantitative RT-PCR. The absolute numbers of viral RNA copies were determined by a standard curve using a viral RNA standard [41].

Compared to the mock control, i.m. immunization with mRNA-N induced a modest but significant control of MA-SARS-CoV-2 in the mouse lung (mean viral copies for mock vs. vaccine: 7,773,344 vs. 846,359; ∼9-fold reduction) (p<0.0001) (**Fig. 2a**). We also evaluated the protective effect of mRNA-N vaccine in mice following intranasal (i.n.) immunization.

**Figure 2.**
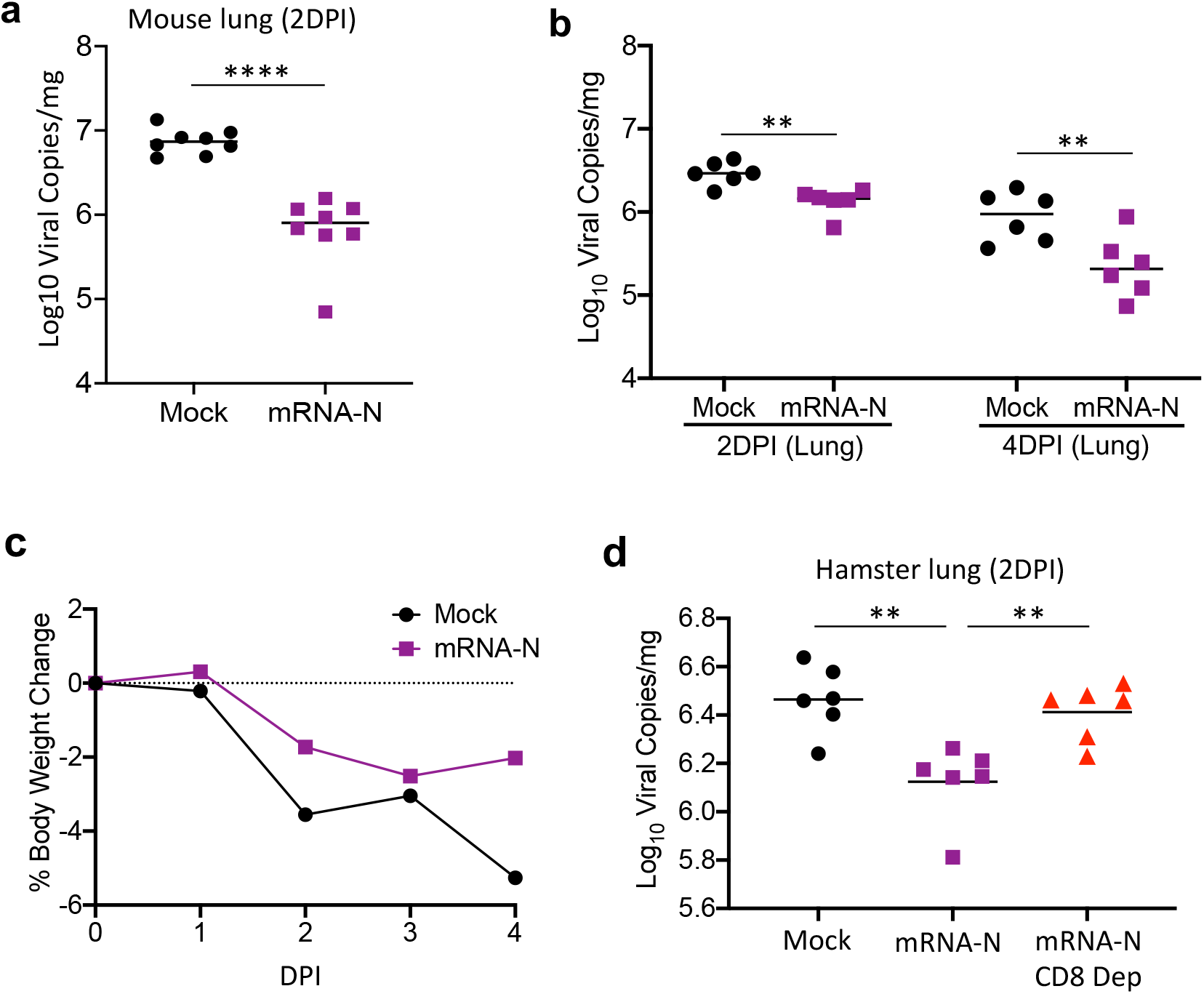
mRNA-N induced protection in mice and hamsters. **(a)** SARS-CoV-2 viral RNA copies (Log_10_) in the lung of mice after vaccination and challenge. Balb/c mice (8/group) were vaccinated with mock and mRNA-N at week 0 and week 3, followed by intranasal challenge with a mouse-adapted SARS-CoV-2 strain (2×10^4^ pfu). Two days post-infection (2 DPI), absolute viral RNA copies in the lung were quantified by qRT-PCR that included a standard curve and were compared between the mock and vaccine group. **(b)** SARS-CoV-2 viral RNA copies (Log_10_) in the lung of hamsters after vaccination and challenge. Hamsters (12/group) were vaccinated with mock and mRNA-N at week 0 and week 3, followed by intranasal challenge with the SARS-CoV-2 delta strain (2×10^4^ pfu). On 2 DPI (n=6) and 4 DPI (n=6), absolute viral RNA copies in the lung were quantified and compared between the mock and vaccine group. **(c)** Comparison of weight loss for hamsters between mock and vaccine group from Day 0 to 4 DPI. **(d)** SARS-CoV-2 viral RNA copies (Log_10_) in the lung of hamsters after vaccination and challenge. Data for mock, mRNA-N, and mRNA-N with CD8 depletion groups were shown. One-way ANOVA or Student’s t-test were used for statistical analysis. * p<0.05, ** p<0.01, *** p<0.001, **** p<0.0001.

Immunization schedules and vaccine dose were identical to those involving the i.m. route. In contrast to i.m. immunization, i.n. immunization with the mRNA-N vaccine failed to induce viral control in the mouse lung (**Fig. S3b**). Consistently, no antibody response was induced in the sera following mRNA-N i.n. immunization (**Fig. S3c**). Thus, i.m. immunization was employed for all subsequent animal experiments.

Next, we evaluated the effectiveness of the mRNA-N vaccine alone against the SARS-CoV-2 delta strain in hamsters. Three cohorts were investigated (**Fig. S3d**). The first two groups (12/group) were i.m. vaccinated with PBS (mock) or mRNA-N vaccine (2µg), respectively, at week 0 and 3, followed by intranasal challenge with the SARS-CoV-2 delta strain (2×10^4^ pfu). Vaccine dose was selected based on the previous studies evaluating mRNA CoVID19 vaccines in hamsters [42, 43]. For each group, on 2 DPI (n=6) and 4 DPI (n=6), all hamsters were analyzed for viral copies in the lung as well as in the upper respiratory tract (nasal wash). In addition, we included a third group (n=6) that received the same mRNA-N vaccine and viral challenge but underwent *in vivo* CD8-cell depletion three days prior to the viral challenge using a well-characterized depletion antibody [44, 45] (**Fig. S3d**). We found that, compared to the mock control, the mRNA-N vaccine induced a modest but significant control of the delta strain (∼3-fold reduction in viral copies) in the lung of hamsters at both 2 DPI (p<0.01) and 4 DPI p<0.01) (**Fig. 2b**). We noted that, compared to the MA-SARS-CoV-2 in mice, the viral suppressive effect of mRNA-N against the delta strain in hamsters appeared to be weaker.

Hamsters were monitored from the day of viral challenge (D0) to 4 DPI, and it was observed that infection with the delta strain led to significant weight loss in hamsters of the mock-vaccinated group (>5% weight loss on 4 DPI); however, only a trend towards reduced weight loss was observed for the mRNA-N-vaccinated hamsters compared to mock-vaccinated hamsters with significant difference detected between the two groups (**Fig. 2c**). These data indicated a modest protection conferred by mRNA-N vaccination against SARS-CoV-2 delta challenge in hamsters, consistent with the modest effect of the vaccine on viral control in the lung of hamsters shown above (**Fig. 2b**). Of particular interest, CD8 depletion largely abrogated the modest effect of the mRNA-N vaccine on viral control in the lung (mean viral copies/mg for mock vs. mRNA-N vs. mRNA-N/CD8 depletion: 3,036,909 vs. 1,397,343 vs. 2,658,119) (p<0.01 for mRNA-N vs. mRNA-N/CD8 depletion) (**Fig. 2d**), indicating a role of CD8^+^ T cells in mediating the effect of mRNA-N vaccine. Finally, we measured viral RNA copies in the nasal washes and found that, compared to the mock control, mRNA-N vaccination did not reduce the viral copies (**Fig. S3e)**. These results indicated that mRNA-N alone, while inducing modest viral control in the lung, had minimal impact on the virus in the upper respiratory tract, likely due to lack of neutralizing antibody induction after vaccination.

### Combinatorial mRNA vaccination leads to more potent protection against SARS-CoV-2 delta variant

After demonstrating that mRNA-N alone was immunogenic and elicited modest efficacy against SARS-CoV-2 in two different animal models, we next explored whether a bivalent platform consisting of mRNA-N with the approved S-expressing mRNA vaccine (mRNA-S-2P) would induce more robust protection. Similarly, combinatorial mRNA-S+N vaccination as compared to the mRNA-S alone was evaluated in both mouse and hamster models. First, three groups of WT Balb/c mice (8/group) were immunized with either PBS (mock), mRNA-S (1µg), or combinatorial mRNA-S & mRNA-N (1µg for each) (mRNA-S+N) as described above at week 0 and week 3, followed by intranasal challenge with the mouse-adapted SARS-CoV-2 strain (2×10^4 pfu) at week 5 (**Fig. S4a**). On 2 DPI, absolute numbers of viral RNA copies in the mouse lung were measured by qRT-PCR as described above. Compared to the mock-vaccinated control, mRNA-S alone was highly effective in controlling the MA-SARS-CoV-2 infection in mice, with no detectable virus in 1 out of 8 mice and weakly detectable viral copies in 7 out of 8 mice (mean viral copies/mg for mock vs. mRNA-S: 7,773,344 vs. 341; >22,000-fold reduction; p<0.0001) (**Fig. 3a**). These data indicate that, unlike mRNA-N, mRNA-S itself is highly efficacious against MA-SARS-CoV-2. Importantly, combinatorial mRNA-S+N vaccination induced an even more robust viral control in the mouse lung, leading to complete viral control with no detectable virus by p-PCR in all 8 mice (p<0.001 for mRNA-S vs. mRNA-S+N) (**Fig. 3a**).

**Figure 3.**
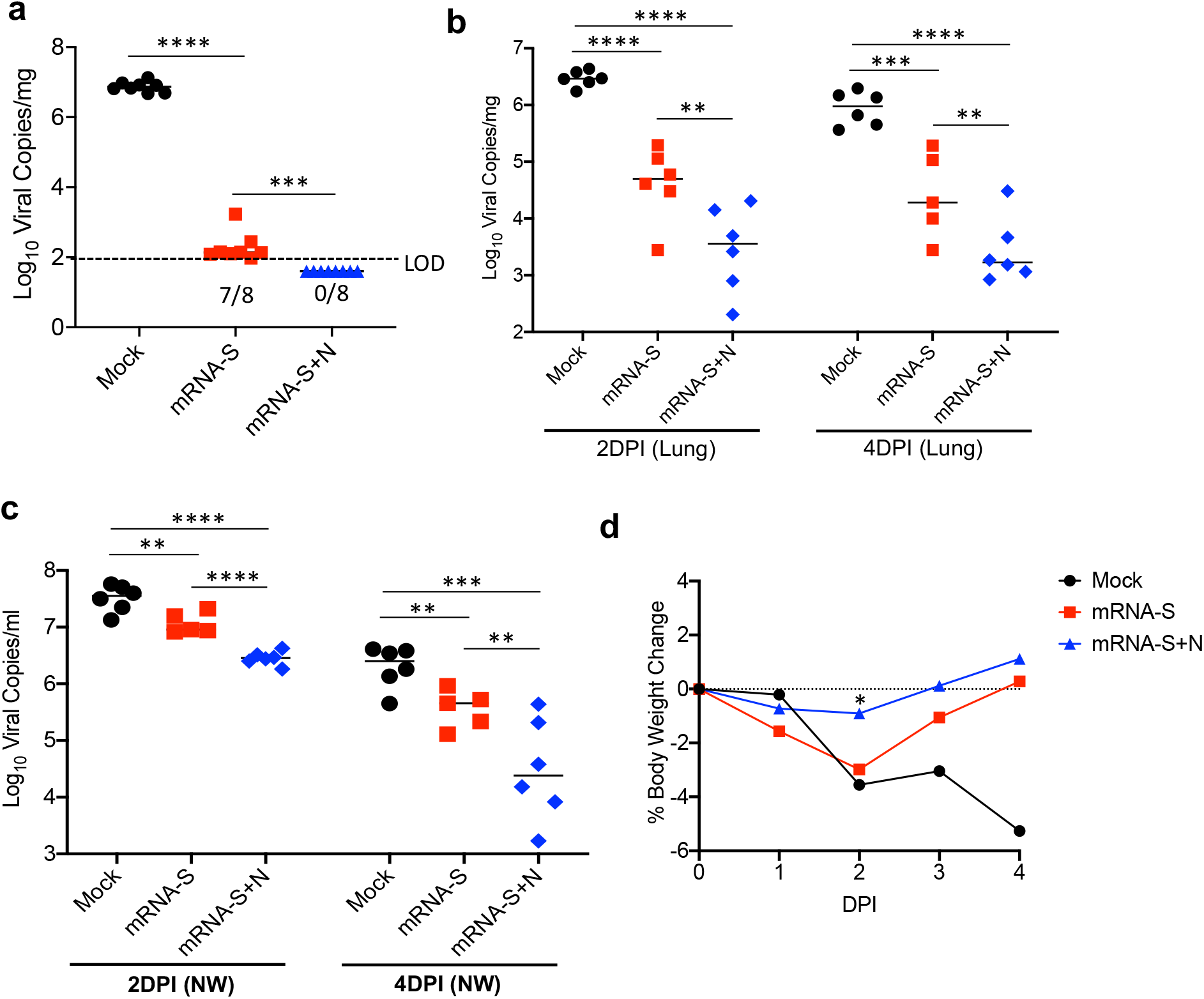
Protection induced by combinatorial mRNA-S+N vaccination compared to mRNA-S alone. **(a)** Viral RNA copies (Log_10_) in the lung of mice after vaccination and challenge. Balb/c mice (8/group) were vaccinated with mock, mRNA-S alone, and combinatorial mRNA-S+N at week 0 and week 3, followed by intranasal challenge with a mouse-adapted SARS-CoV-2 strain (2×10^4^ pfu). On 2 DPI, absolute viral RNA copies in the lung were quantified. **(b-c)** Viral RNA copies (Log_10_) in the lung **(b)** or nasal wash **(c)** of hamsters after vaccination and challenge. Hamsters (12/group) were vaccinated with mock, mRNA-S alone, and combinatorial mRNA-S+N at week 0 and week 3, followed by intranasal challenge with the delta strain (2×10^4^ pfu). On 2 DPI (n=6) and 4 DPI (n=6), absolute viral RNA copies in the lung (b) or in nasal wash (d) were quantified. **(d)** Comparison of weight loss for hamsters between mock and vaccine groups from Day 0 to Day 4 after viral infection (DPI). One-way ANOVA was used for statistical analysis. * p<0.05, ** p<0.01, *** p<0.001, **** p<0.0001.

Second, to more vigorously evaluate the effect of combinatorial mRNA-S+N vaccination as compared to mRNA-S alone, we employed the hamster model as described above. Three groups (12/group) were vaccinated with PBS (mock), mRNA-S (2 µg), or combinatorial mRNA-S & mRNA-N (2 µg for each) at week 0 and 3, followed by intranasal challenge with the SARS-CoV-2 delta strain at week 5 (**Fig. S4b)**. Within each group, animals were monitored for signs of morbidity and weight loss. On 2 DPI (n=6) and 4 DPI (n=6), hamsters were analyzed for vaccine-induced protection based on body weight loss and viral RNA copies in the lung as well as in nasal washes. On 2 DPI, compared to the mock-vaccinated control, mRNA-S alone induced substantial reduction in viral copies in the lung (mean copies/mg for mock vs. mRNA-S: 3,036,906 vs. 73,578; ∼45-fold reduction; p<0.0001) (**Fig.3b**), which is consistent with clinical observations that mRNA-S vaccine is effective against disease caused by the delta variant [46]. However, relative to the MA-SARS-CoV-2 infection in mice (almost complete viral control by mRNA-S), mRNA-S alone was less effective in controlling the delta variant in hamsters. Critically, compared to mRNA-S alone, combinatorial mRNA-S+N vaccination induced more robust control of the virus by inducing an additional ∼10-fold reduction of viral copies in the lung on 2 DPI (mean viral copies/mg for mRNA-S vs. mRNA-S+N: 73,578 vs. 7178; p<0.01) (**Fig. 3b**). Compared to the mock-vaccinated control, there was ∼450-fold reduction in viral copies by combinatorial mRNA-S+N vaccination (p<0.0001) (**Fig. 3b**). A similar result was observed on 4 DPI, where compared to the mock-vaccinated control, the reduction in viral copies induced by mRNA-S or mRNA-S+N was 16-fold and 160-fold, respectively (**Fig. 3b**).

Viral loads in the upper respiratory tract have implications for SARS-CoV-2 transmission and potential breakthrough infections. Therefore, viral RNA copies in nasal washes were also quantified in these hamsters on 2 DPI and 4 DPI. As shown in **Fig. 3c**, unlike the robust viral control by mRNA-S in the lung, mRNA-S alone was less effective in reducing viral copies in the nasal wash on both 2 DPI (mean viral copies/ml for mock vs. mRNA-S: 35,968,374 vs. 12,562,995; ∼3-fold; p<0.01) and 4 DPI (mock vs. mRNA-S: 2,505,226 vs. 450,804; ∼6-fold; p<0.01) (**Fig. 3c**). These data indicate a possible explanation that the mRNA-S vaccine tends to provide strong protection against disease but reduced protection against infection caused by the delta variant [29]. Notably, compared to mRNA-S alone, combinatorial mRNA-S+N vaccination induced stronger viral control in the nasal washes on 2 DPI (∼4-fold reduction in viral copies relative to mRNA-S alone, p<0.0001; ∼12-fold reduction relative to the mock control, p<0.0001) and 4 DPI (∼4-fold reduction relative to mRNA-S alone, p<0.05; ∼21-fold reduction relative to mock control, p<0.001) (**Fig. 3c**). Together, these data support that combinatorial mRNA-S+N vaccination induces stronger and faster control of the SARS-CoV-2 delta infection in both lung and upper respiratory tract, indicating that this vaccine approach may also reduce the risk of viral spread and transmission.

Analysis of hamster body weights showed that challenge with the delta variant caused progressive weight losses in the mock group, declining by >5% on 4 DPI (**Fig. 3d**), which is comparable with that caused by the WT SARS-CoV-2 [41]. Compared to the mock-vaccinated control, vaccination with mRNA-S alone or mRNA-S+N both prevented hamsters from weight loss on 3 DPI and 4 DPI (**Fig. 3d**). Compared to mRNA-S alone, there was a trend towards better protection by mRNA-S+N with a significant difference detected on 2DPI (p<0.05) (**Fig. 3d**).

### Combinatorial mRNA vaccination elicits robust N- and S-specific T-cell and humoral immunity

We next sought to understand immune parameters associated with the enhanced protection by combinatorial mRNA vaccination as compared to mRNA-S alone or mock-vaccinated control. A mouse immunogenicity study was conducted, where three groups of Balb/c mice (7/group) were vaccinated with PBS (mock), mRNA-S alone, or mRNA-S+N combination at week 0 (prime) and week 3 (boost) using a similar experimental design as described in **Fig. S4a**, except that at week 5 (2 weeks after booster) all mice were subjected to immune analysis. Flow cytometric analysis of total T-cell activation (based on CD44+) showed that compared to the un-vaccinated control, both mRNA-S and mRNA-S+N elicited strong activation of CD4^+^ and CD8^+^ T cells in mice, which did not differ significantly between the mRNA-S and mRNA-S+N groups (**Fig. 4a**). Splenocytes were re-stimulated with S or N peptide pools and ICS was performed for identifying vaccine-induced, S- and N-specific T cells based on cytokine expression (IFN-γ, TNF-α, IL-2) (**Fig. 4b-e**). The data showed that combinatorial vaccination elicited robust S-specific (**Fig. 4b-c**) as well as N-specific (**Fig. d-e**) CD4^+^ and CD8^+^ T-cell responses in mice. Among the cytokines examined, TNF-α was highly expressed by both S- and N-specific T cells, followed by IFN-γ and IL-2 (**Fig. 4b-e**). When comparing with mRNA-S alone, combinatorial mRNA-S+N vaccination appears to induce some synergistic effects and augments the S-specific CD8^+^ T-cell response (TNF-α+ or IFN-γ+) (**Fig. 4c)**. As a control, very little or no N-specific T-cell response was detected in the mRNA-S alone group (**Fig. 4d-e)**. Induction of both S- and N-specific T-cell responses by combinatorial mRNA-S+N vaccination as compared to the mRNA-S alone was further confirmed by IFN-γ ELISPOT (**Fig. S5a-b**).

**Figure 4.**
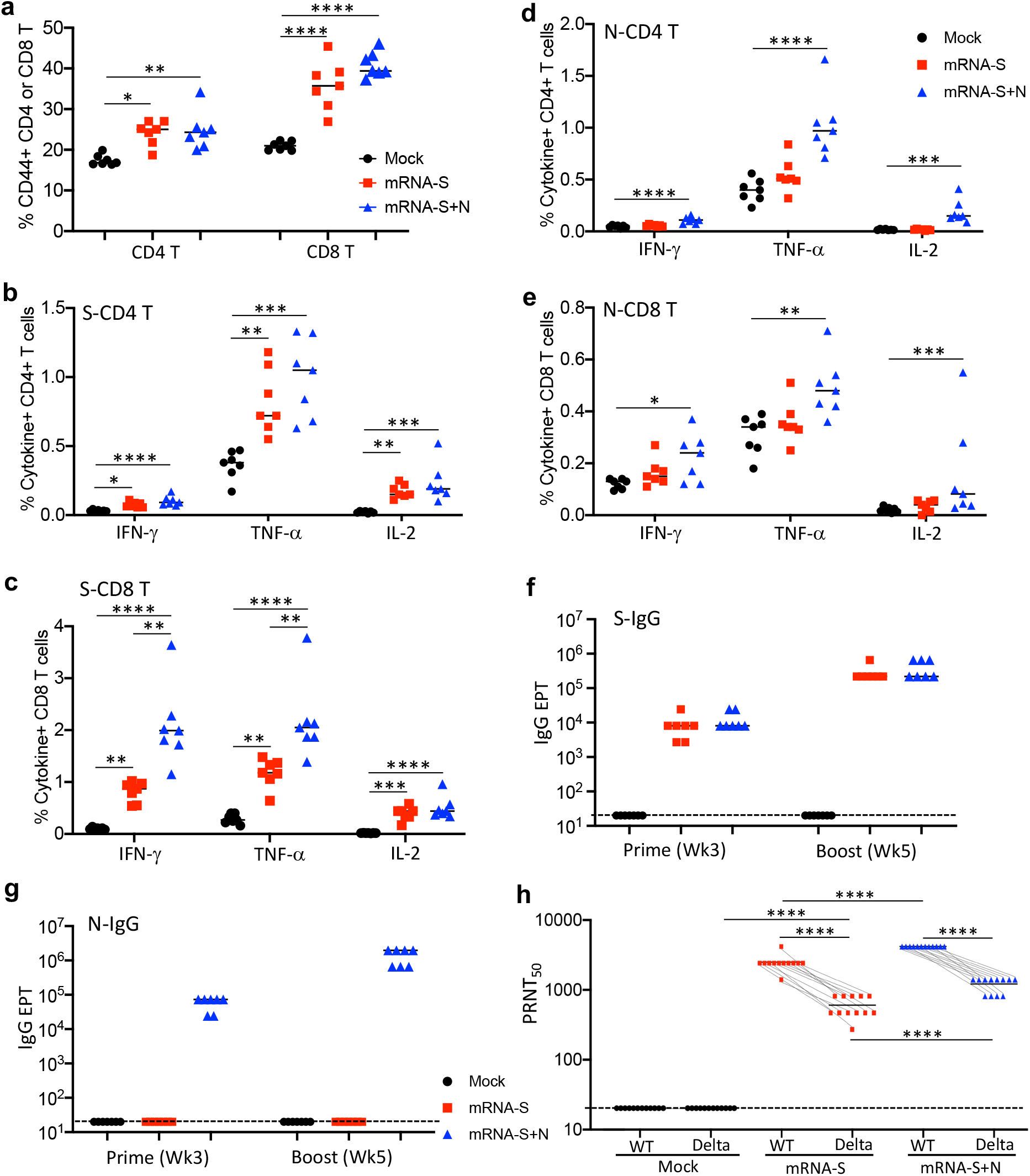
Immune analysis of mRNA-S and combinatorial mRNA-S+N vaccination in mice and hamsters. **(a)** Analysis of total CD4^+^ and CD8^+^ T cell activation in the mouse spleen following vaccination. Three groups of mice (7/group) were vaccinated with mock, mRNA-S, or combinatorial mRNA-S+N as indicated above. Splenocytes collected 2 weeks after booster were stained and the expression of CD44 on CD4^+^ and CD8^+^ T cells were examined by flow cytometry and shown as % CD44^+^ in CD4^+^ or CD8^+^ T cells. **(b-c)** ICS measurement of S-specific CD4^+^ and CD8^+^ T cells in the mouse spleen. % individual cytokine-positive, S-specific CD4^+^ (b) or CD8^+^ (c) T cells were examined and compared between mock and vaccine groups. (**d-e**) ICS measurement of N-specific CD4^+^ and CD8^+^ T cells in the mouse spleen. % individual cytokine-positive, N-specific CD4^+^ (d) or CD8^+^ (e) T cells were examined and compared between mock and vaccine groups. **(f-g)** ELISA measurement of serum S-specific **(f)** or N-specific **(g)** binding IgG following prime (week 3) or booster (week 5) vaccination in mice. Antibody endpoint titers (EPT) were determined based on serum serial dilutions (1:3 ratio) and were shown for the mock and vaccine groups after prime and booster vaccination. **(h)** Hamster serum neutralizing activity against WT SARS-CoV-2 and the delta variant. Serum samples collected from the hamsters (Fig. 3b) after booster vaccination (week 5) but prior to viral challenge were measured for neutralizing activity by PRNT. PRNT_50_ for individual serum samples of each group were shown and compared among different groups as well as between the WT virus and the delta variant within each group. Dotted line in each plot showed limit of detection for each assay. One-way ANOVA or Student’s t-test were used for statistical analysis. * p<0.05, ** p<0.01, *** p<0.001, **** p<0.0001.

Serum binding IgG specific to S or N proteins in these mice were also analyzed by ELISA and the data revealed similar patterns (**Fig. 4f-g**). As expected, mRNA-S alone induced robust binding IgG specific to S (**Fig. 4f**), but not to N (**Fig. 4g**), following prime vaccination (mean S-IgG EPT: 8,871), which was markedly enhanced by booster vaccination (mean S-IgG EPT: 281,186) (**Fig. 4f**). Compared to mRNA-S alone, combinatorial mRNA-S+N elicited strong binding IgG specific to both S and N proteins following prime vaccination (mean S-IgG EPT: 12,729; mean N-IgG EPT: 59,014), both of which were also enhanced by booster vaccination (mean S-IgG EPT: 406,157; mean N-IgG EPT: 1,405,929) (**Fig. 4f-g)**. Of note, compared to mRNA-S alone, combinatorial mRNA-S+N vaccination also appeared to provide some synergistic effects and modestly augmented the S-specific binding IgG, although no statistical significance was detected (**Fig. 4f**).

We next evaluated vaccine-induced serum neutralizing activities. In the hamster experiments described earlier (**Fig. 3b-c)**, serum samples were collected after booster vaccination (week 5) and prior to SARS-CoV-2 challenge. Their neutralizing activities against the delta variant and the WT SARS-CoV-2 (WA1/2020) was measured in parallel by the PRNT assay [47]. Sera of the mRNA-S vaccinated hamsters manifested strong neutralizing activity against WT virus (mean WT PRNT_50_: 2667), whereas their neutralizing activities against the delta variant was markedly reduced (mean delta PRNT_50_: 440; ∼5-fold reduction) (**Fig. 4h**), although mRNA-S-induced delta neutralizing activity remained significantly higher compared to the mock control (mean delta PRNT_50_ for mock vs. mRNA-S: <20 vs. 440) (**Fig. 4h**). Of importance, compared to mRNA-S alone, combinatorial mRNA-S+N vaccination elicited stronger serum neutralizing activity against the delta variant (mean delta PRNT_50_ for mRNA-S vs. mRNA-S+N: 440 vs. 1067; increase by ∼1.5 fold) (p<0.001) as well as against the WT virus (mean WT PRNT_50_ for mRNA-S vs. mRNA-S+N: 2667 vs. 5120; increase by ∼0.9 fold) (**Fig. 4h**). These data were consistent with the augmented S-specific CD8 T-cell response (**Fig. 4c**) by mRNA-S+N vaccination compared to mRNA-S alone and indicated that presence of N antigen for immunization likely induces an immune environment that promotes the generation of S-specific immunity (to be discussed subsequently in Discussion). Together, our immune analyses suggest that combinatorial mRNA-S+N vaccination not only induces N-specific immunity, but also elicits a stronger S-specific CD8 T-cell response and serum neutralizing antibody activity when compared to mRNA-S alone, which may collectively contribute to the enhanced protection against SARS-CoV-2 delta variant infection.

## DISCUSSION

We report a new nucleoside-modified mRNA vaccine that expresses the SARS-CoV-2 nucleoprotein (mRNA-N) and its immunogenicity and protective efficacy when used alone or in combination with the current approved, S-expressing mRNA vaccine (mRNA-S-2P). We demonstrate that mRNA-N is highly immunogenic and by itself induces modest but significant control of SARS-CoV-2 in mice and hamsters. Additionally, compared to the mRNA-S alone, combinatorial mRNA-S+N vaccination provides markedly stronger protective efficacy against SARS-CoV-2 delta variant in the lung (by additional 10-fold reduction in viral copies compared to mRNA-S and ∼450-fold reduction compared to the mock-vaccinated control) and in the upper respiratory tract. Immune analyses indicate that induction of N-specific immunity together with the augmented S-specific immunity likely contribute to the enhanced host protection conferred by the combinatorial mRNA vaccination. Thus, our study reports a new mRNA vaccine approach for improved control of the circulating delta variant and thus suggests strategies for the development of next-generation vaccines for SARS-CoV-2 variants (e.g., omicron) and pan-coronaviruses in the near future.

The majority of current CoVID19 vaccines principally targets the viral S protein (reviewed in [22, 48, 49]). Emergence of SARS-CoV-2 VOCs, including the highly transmissible delta strain and newly emerged omicron, has posed challenges to the currently approved vaccines [24-28]. Compared to the S protein, less is known about the role of the viral N protein in SARS-CoV-2 vaccination. In this study, we generated pseudouridine-modified, LNP-formulated mRNA that encodes the full-length SARS-CoV-2 N protein and demonstrated that mRNA-N is highly immunogenic, and able to elicit N-specific T cell response (both CD4 and CD8) (**Fig. 1a-e**) and strong binding IgG response (**Fig. 1f-g**). As expected, no serum neutralizing activity was induced by mRNA-N (**Fig. 1h**). Thus, the mRNA-N vaccine provides an opportunity to explore whether the immune response to the N protein alone could confer protection against SARS-CoV-2 in the absence of neutralizing antibodies. Two different animal models, including WT mice infected with mouse-adapted SARS-CoV-2 [40] and hamsters infected with the delta strain (**Fig. 2**), were employed in our study and both models showed that mRNA-N immunization (intramuscular) induced a modest but significant control of SARS-CoV-2 in the lung (∼10-fold reduction in mice and ∼3-fold reduction in hamsters). However, relative to mRNA-S, which induced almost complete control of mouse-adapted SARS-CoV-2 in mice (**Fig. 3a**) and robust control of the delta strain (by 45-fold) in hamsters (**Fig. 3b**), the viral control by mRNA-N vaccine alone was only modest (**Fig. 2a-b**), supporting a major role for the S protein in immune protection against SARS-CoV-2. We also explored mucosal delivery of mRNA-N and observed that neither serum antibody response nor viral control was observed following mRNA-N intranasal vaccination (**Fig. S3b-c)**, indicating that it i.n. delivery may not be a favorable route for mRNA vaccination.

With vaccination rates increasing and constant mutations of viral S proteins, multiple variants have been identified that showed reduced sensitivity to vaccine-induced neutralization [26-28]. Clinical studies also indicate that the two approved mRNA-S vaccines (BNT162b2 and mRNA-1273), while retaining high efficacy in protecting from hospitalization and death caused by the delta variant, have limited overall efficacy in preventing infection, which was greatly reduced (BNT162b2: 51.9% (95% CI, 47.0–56.4%); mRNA-1273: 73.1% (95% CI, 67.5–77.8%); both for ≥14 d after the second dose) [29]. In agreement with these findings, our data showed that while the mRNA-S vaccine was highly effective against a mouse-adapted SARS-CoV-2 strain and almost completely controlled the virus (**Fig. 3a**), its efficacy was reduced against the delta strain in hamsters (**Fig. 3b**). Notably, the protective effect of the mRNA-S vaccine from the delta strain appeared to be even weaker in the upper respiratory tract (in nasal washes, ∼3 to 5-fold reductions in viral copies compared to the mock-vaccinated control on both 2 and 4 DPI) (**Fig. 3c**). These data provide a possible explanation why mRNA-S vaccination showed reduced efficacy against delta infection in clinical studies. A caveat of the present study is lack of WT SARS-CoV-2 challenge controls in the hamster experiments, which could be used for direct comparison with the delta challenge. However, based on previous preclinical testing of BNT162b2 and mRNA-1273 using WT virus, both mRNA-S vaccines were found to be highly effective in controlling WT SARS-CoV-2 in both upper and lower respiratory tracts [5, 8]. In addition, our analyses of serum neutralizing activity against WT and delta strains in parallel (**Fig. 4h**) also supported these observations as sera of mRNA-S vaccinated hamsters manifested reduced neutralizing activities by ∼5-fold against the delta strain compared to WT virus.

A major finding of our study is that combinatorial mRNA-S+N vaccination led to significant increase in control of the SARS-CoV-2 delta strain infection in the lungs (**Fig. 3b**) and in the upper respiratory tract (**Fig. 3c**). Notably, combinatorial vaccination induced an additional 10-fold reduction in viral RNA copies compared to the mRNA-S alone (∼450-fold reduction in the lungs compared to the mock-vaccinated control at 2 DPI) and led to a faster resolution of the infection, indicating that this vaccine approach likely provides even better protection from the disease caused by delta. Furthermore, combinatorial vaccination also induced stronger viral control in the upper respiratory tract (by additional 4-fold compared to mRNA-S alone) (**Fig. 3c**), albeit to lesser extent than that in the lung. This finding indicates that combinatorial mRNA-S+N vaccination may also reduce the risk of transmission by delta. Collectively, given the constant mutations of S gene and the generation of SARS-CoV-2 spike variants with partial escape from vaccine-induced immunity [49, 50], our data support that simultaneous targeting of the S protein and another conserved antigen of the virus could confer some broader protection. In future studies, it would be interesting to further explore if this vaccine concept could be harnessed to induce protection against additional SARS-CoV-2 variants (e.g., omicron) and other coronaviruses (e.g., SARS-CoV-1 and MERS).

Another noteworthy observation of our study is that, compared to the mRNA-S alone, combinatorial mRNA-S+N vaccination led to augmented S-specific CD8^+^ T-cell immunity (**Fig. 4c**) and serum neutralizing activities (**Fig. 4h**). This finding is somewhat surprising since the doses of the mRNA-S in the two groups were identical. This is also unlikely due to the extra lipid nanoparticles (which is present in the mRNA-N vaccine) administered in the mRNA-S+N group, because a recent study evaluating various doses of mRNA-1273 vaccine in hamsters showed comparable levels of neutralizing antibodies induced by 1 µg and 5 µg of mRNA vaccines [43]. Instead, our data indicate that mRNA-N immunization may provide some synergistic effects on the generation of S-specific immunity. The mechanisms remain unclear, but we speculate that presence of mRNA-N primes a stronger or more favorable innate immune response, which promotes the generation of S-specific T-cell and humoral immunity. Better understanding of the mechanisms for the coordinated induction and regulation of S- and N-specific immunity following combinatorial mRNA vaccination is needed. For example, a more detailed dissection of early events following combinatorial vaccination, such as protein expression, antigen presentation, and stimulation of innate immune responses, would be helpful to address these questions. In addition, the current study revealed multiple key differences in antigen-specific adaptive immunity elicited by combinatorial mRNA vaccination compared to vaccination with mRNA-S alone. These included the induction of N-specific T-cell response (**Fig. 4d-e**) and the augmented S-specific T-cell response (**Fig. 4c**) and neutralizing activities (**Fig. 4h**), which may collectively contribute to the enhanced immune protection conferred by combinatorial mRNA vaccination. However, the role of these individual immune responses in the enhanced vaccine efficacy remains unclear and should be further explore. This knowledge is thought to not only help understand host protective immunity to coronavirus infection, but also inform strategies for design of pan-coronavirus vaccines.

Durability of vaccine-induced immunity is another critical issue [22]. Studies on coronavirus-infected patients indicated that viral infection can induce immunological memory ranging from months to years, but long-term data on SARS-CoV-2-induced immunity remains lacking [50]. Monitoring immune responses in the COVID-19 mRNA vaccinated individuals indicates that vaccine-induced antibody response wanes over time [51, 52]. Some previous studies on host immunity to coronaviruses (e.g., SARS-CoV-1 and MERS) showed that T-cell immunity could be maintained for longer periods of time compared to antibody responses [53, 54]. While we demonstrated here that combinatorial vaccination induced robust T-cell and humoral immunity specific to both S and N antigens, durability of these immune responses remains unclear, which represents a limitation of the current study. Additional experiments are warranted to determine the durability of immunity and potential long-term protection conferred by the combinatorial mRNA vaccination compared to mRNA-S alone in future research.

In summary, our data demonstrate that combinatorial mRNA vaccination markedly enhances viral control of SARS-CoV-2 including the delta variant, providing a proof-of-concept that a vaccine approach targeting both viral S protein and additional conserved region of the virus is feasible and could induce stronger and broader protection against VOCs. This vaccine approach has potential to facilitate the control of COVID-19 pandemic and warrants further investigation and development.

## METHODS

### mRNA synthesis and LNP formulation

Antigens encoded by the mRNA vaccines in this study were derived from SARS-CoV-2 isolate Wuhan-Hu-1 (GenBank MN908947.3). Nucleoside modified mRNAs expressing SARS-CoV-2 full-length N (mRNA-N) or full-length S with two proline mutations (mRNA-S-2P) were synthesized by in vitro transcription using T7 RNA polymerase (MegaScript, Ambion) on linearized plasmid templates as previously reported [35]. UTP was replaced with One-methylpseudouridine (m1Ψ)-5’-triphosphate (TriLink, Cat# N-1081) for producing nucleoside-modified mRNAs. Poly-A tail was added to the end of modified mRNAs for optimized protein expression. *In vitro* transcribed mRNAs were capped using ScriptCap m7G capping system and ScriptCap 2′-O-methyl-transferase kit (ScriptCap, CellScript) [35], followed by purification using cellulose purification method [36]. Purified mRNAs were analyzed by agarose gel electrophoresis and were kept frozen at -20°C. mRNAs were formulated into lipid nanoparticles (LNP) using an ethanolic lipid mixture of ionizable cationic lipid and an aqueous buffer system as previously reported [37, 55]. Formulated mRNA-LNPs were prepared according to RNA concentrations (1 µg/µl) and were stored at -80°C for animal immunizations.

### Western blot analysis of protein expression by mRNA-N

293T cells in 6-well plate were directly transfected with 2 µg of mRNA-N-LNP or not transfected (as a cell-only control). 18 hours after transfection, cells were lysed in RIPA buffer (Thermo Fisher Scientific) for western blot (WB) analysis. Cell lysates were centrifuged, followed by collection of supernatants for quantification of total protein concentration using Microplate BCA Protein Assay Kit (Pierce™, Thermo Fisher Scientific). Equal amounts of protein were separated by SDS-PAGE using 4-15% SDS polyacrylamide gels (Bio-Rad). Proteins were then transferred onto a nitrocellulose membrane (Bio-Rad). The membrane was blocked in tris buffered saline (TBS) containing 0.05% Tween-20 (Thermo Fisher Scientific) and 5% (w/v) non-fat dried milk (Bio-Rad) for 1 hour at room temperature, followed by incubation with anti-SARS-CoV2 nucleocapsid mouse mAb (MA5-29981, Invitrogen; 1:1000) overnight at 4°C. After washing in TBST (3 times for 5 min), the membrane was incubated for 1 hour with HRP-linked anti-mouse IgG (7076S, Cell Signaling; 1:5000). The membrane was washed, and proteins were visualized using the ECL Western Blotting Substrate (Thermo Fisher Scientific).

### Animal ethics statement

The animal study protocols were approved by the Institutional Animal Care and Use Committee (IACUC) at the University of Texas Medical Branch. Animal studies were conducted in accordance with the recommendations in the Guide for the Care and Use of Laboratory Animals of the National Institutes of Health.

### Mouse immunization and SARS-CoV-2 challenge

Vaccine immunogenicity and effectiveness of protection were evaluated in WT Balb/c mice. 6-week-old female BALB/c mice were obtained from the Jackson Laboratories (Wilmington, MA, USA) and were housed in the animal facility of the University of Texas Medical Branch. For immunogenicity analysis, four groups of mice (7/group) were immunized intramuscularly (i.m.) with either PBS (mock control), mRNA-S (1 µg), mRNA-N (1 µg), or combined mRNA-S+N (1 µg for each) using a prime-boost approach at week 0 (prime) and week 3 (boost), respectively. The vaccine or control was administered at 50 µl per injection. Blood/serum samples were collected from all mice three weeks after prime vaccination (prior to booster vaccination) for measuring vaccine-induced antibody response. Two weeks after booster vaccination (week 5), all mice were euthanized. Blood/serum and spleen samples were collected for analyses of vaccine-induced humoral and cellular immune responses.

For mouse challenge study, another four groups of BALB/c mice (8/group) received the same mock control or vaccines as indicated above. Vaccine doses and immunization timeline were identical to the above immunogenicity study. Two weeks after booster vaccination (week 5), all mice were transferred to the ABSL-3 facility and were intranasally challenged with a mouse-adapted SARS-CoV2 CMA4 strain (2×10^4^ pfu) as previously reported [40]. The details of this mouse model were recently published [56]. Two days after viral challenge, all mice were euthanized, and equivalent portions of the lung tissues were collected for quantification of SARS-CoV-2 viral copies.

### Hamster immunization and SARS-CoV-2 delta strain challenge

Vaccine-induced protection against the SARS-CoV-2 delta strain was evaluated in hamsters. Four groups of 4- to 5-week-old male golden Syrian hamsters (12/group), strain HsdHan: AURA (Envigo, Indianapolis, IN), were vaccinated intramuscularly (i.m.) with either PBS (mock control), mRNA-S (2 µg), mRNA-N (2 µg), or combined mRNA-S+N (2 µg for each) using a prime-boost approach at week 0 (prime) and week 3 (boost), respectively. The vaccine or control was administered at 100 µl per injection. Blood/serum samples were collected from all mice three weeks after prime vaccination (week 3) and two weeks after booster vaccination (week 5) for measuring vaccine-induced neutralizing activities. 14 days after booster vaccination (week 5), all hamsters were transferred to the ABSL-3 facility and were intranasally challenged with the SARS-CoV2 delta strain (2×10^4^ pfu) (World Reference Center for Emerging Viruses and Arboviruses). Two days post-infection (2 DPI), 6 hamsters of each group were euthanized. Nasal wash samples and equivalent portions of the lung tissues were collected from these hamsters for quantification of SARS-CoV-2 viral loads in the upper respiratory tract and in the lung, respectively. Four days post infection (4 DPI), the same procedures were conducted for the other 6 hamsters in each group. Body weights of hamsters were monitored from the day of viral challenge to day 4 post viral challenge (0 DPI to 4 DPI) to evaluate vaccine-induced protection from animal weight loss.

### Binding IgG by ELISA

ELISA was conducted to measure vaccine-induced, N- and S-specific binding IgG in the mouse sera. ELISA plates (Greiner bio-one) were coated with 1µg/ml recombinant S (S1+S2-ECD; 40589-V08B1; Sino Biological) or N protein (40588-V08B; Sino Biological) in DPBS overnight at 4°C. Plates were washed three times with wash buffer (DPBS with 0.05% Tween 20), 5 min each time, and then blocked with 8% FBS in DPBS for 1.5 hour at 37°C. Plates were washed and incubated with serially diluted sera (initial dilution 1:100; 1:3 serial dilution) in blocking buffer at 50 µl per well for 1 hour at 37°C. Plates were washed again and then incubated with horse radish peroxidase (HRP) conjugated anti-mouse IgG secondary antibody (Biolegend; 1:3000) for 1 hour at 37°C. After the final wash, plates were developed using TMB 1-Component Peroxidase Substrate (Thermo Fisher), followed by termination of reaction using the TMB stop solution (Thermo Fisher). Plates were read at 450 nm wavelength within 15 min by using a Microplate Reader (BioTek). Binding IgG Endpoint titers (EPT) for each sample were calculated.

### Neutralizing assay

Serum neutralizing activity was examined by a standard Plaque Reduction Neutralization Test (PRNT) as previously reported [57, 58]. The assays were performed with Vero cells using the SARS-CoV-2 WT or delta strains at BSL-3. In brief, sera were heat-inactivated and two-fold serially diluted (initial dilution 1:10), followed by incubation with 100 PFU WT SARS-CoV2 (USA-WA1/2020) or the delta strain for 1 hour at 37°C. The serum-virus mixtures were placed onto Vero E6 cell monolayer in 6-well plates for incubation for 1 hour at 37°C, followed by addition of 2 ml overlay consisting of MEM with 1.6% agarose, 2% FBS and 1% penicillin–streptomycin to the cell monolayer. Cells were then incubated for 48 hours at 37°C, followed by staining with 0.03% liquid neutral red for 3-4 hours. Plaque numbers were counted and PRNT_50_ were calculated. Each serum sample was tested in duplicates.

### Intracellular Cytokine Staining (ICS) and Flow Cytometry

ICS was performed for splenocytes to detect vaccine-specific T-cell response. Cells were washed with FACS buffer (1% FBS and 0.5 M EDTA in PBS) and resuspended with complete RPMI with 10 mM HEPES supplemented with 10% FBS, 2-Mercaptoethanol, Sodium Pyruvate, Non-Essential Amino Acids, Pen-Strep, and L-Glutamine. Cells were then stimulated with 1 µg/ml S peptide pool (JPT, PM-WCPV-S) or N peptide pool (Meltenyi, 130-126-698) in the presence of 1 µg/mL anti-CD28 (Invitrogen, 14-0281-86) for co-stimulation for 6 hours. In the last 4 hours of incubation, protein transport inhibitor Brefeldin-A was added. Cells stimulated with PMA/Ionomycin or DMSO only were included as positive control and negative control, respectively. Following stimulation, cells were first stained for surface markers, including CD4-percp/cy5.5 (Biolegend, 100540), CD8-BV711 (Biolegend, 100759), and CD44-BV510 (Biolegend, 103044). The surface staining was performed on ice for 30 min. After washing with PBS, cells were resuspended with Zombie-dye (Biolegend) for viability staining and incubated at room temperature (RT) for 15 min. Following surface and viability staining, cells were fixed with fixation buffer (Biolegend, 420801) and permeabilized with perm/wash buffer (Biolegend, 421002), followed by intracellular cytokine staining with IFN-γ-BV605 (Biolegend, 505840), TNF-α-PE/cy7 (Biolegend, 506324), and IL-2-APC (Tonbo bioscience, 20-7021) on ice for 30 min. Cells were then washed with perm/wash buffer and were processed with a multi-parametric flow cytometer FACS LSR Fortessa (BD). Data were analyzed using FlowJo (TreeStar).

### IFN-γ ELISPOT

Millipore ELISPOT plates (Millipore Ltd, Darmstadt, Germany) were coated with anti-IFN-γ capture Ab (CTL, Cleveland, OH, USA) at 4°C overnight. Splenocytes (0.25 × 10^6^) were stimulated in duplicates with SARS-CoV-2 S- or N-peptide pool (2 µg/ml, Miltenyi Biotec, USA) for 24 hours at 37°C. Splenocytes stimulated with anti-CD3 (1 µg/ml, e-Biosciences) or medium alone were used as positive and negative controls, respectively. This was followed by incubation with biotin-conjugated anti-IFN-γ (CTL, Cleveland, OH, USA) for 2 hours at room temperature, and then alkaline phosphatase-conjugated streptavidin for 30 minutes. The plates were washed and scanned using an ImmunoSpot 4.0 analyzer and the spots were counted with ImmunoSpot software (Cellular Technology Ltd, Cleveland, OH) to determine the spot-forming cells (SFC) per 10^6^ splenocytes.

### RNA extraction and qPCR quantification of viral loads

RNA was extracted from the lung tissues (mice and hamsters) and nasal wash (hamsters) using the TRIzol LS reagent (Thermo Fisher Scientific) according to the manufacturer’s instructions. Concentration and purity of the extracted RNAs were determined using the multi-mode plate reader (BioTek). To quantify SARS-CoV-2 viral RNA copies, quantitative RT-PCR was performed using the iTaq Universal SYBR Green One-Step Kit (Bio-Rad) and the CFX Connect Real-Time PCR Detection System (Bio-Rad). Primer set for the SARS-CoV-2 E gene (F: 5’-GGAAGAGACAGGTACGTTAATA-3’; R: 5’-

AGCAGTACGCACACAATCGAA-3’). PCR reactions (20 μl) contained primers (10μM), RNA sample (2 μl), iTaq universal SYBR Green reaction mix (2X) (10 μl), iScript reverse transcriptase (0.25 μl), and molecular grade water. PCR cycling conditions were: 95°C for 3 minutes, 45 cycles of 95°C for 5 seconds, and 60°C for 30 seconds. For each PCR, a standard curve was included, using an RNA standard (*in vitro* transcribed 3,839bp containing genomic nucleotide positions 26,044 to 29,883 of SARS-CoV-2 genome), to quantify the absolute copies of viral RNA in the lung tissue or nasal wash.

### Statistical Analysis

Statistical analyses were performed using Graph-Pad Prism 8.0. Statistical comparison was performed using either unpaired Student’s t test or one-way ANOVA where appropriate. The values were presented either as mean or mean ± SD. Two-tailed p values were denoted, and p values < 0.05 were considered significant.

## ACKNOWLEDGEMENT

The research was supported by a UTMB Institute for Human Infections and Immunity CoVID-19 grant (to H.H), the Sealy and Smith Foundation, and grant R24AI120942 from NIH (to S.C.W.). H.H. was supported by NIH grants AI157852 and AI147903. Y.L. and L.S. were supported by NIH grants AI132674 and AI56536. P.-Y.S. was supported by NIH grants HHSN272201600013C, AI134907, AI145617, and UL1TR001439, and awards from the Sealy & Smith Foundation, the Kleberg Foundation, the John S. Dunn Foundation, the Amon G. Carter Foundation, the Gilson Longenbaugh Foundation, and the Summerfield Robert Foundation. T.W. was supported by NIH grants R01AI127744 and R21 AI140569. R.H. was supported by a UTMB Sealy Institute for Vaccine Sciences Fellowship and the McLaughlin Fellowship. We thank UTMB’s flow cytometry and cell sorting core for assistance in flow cytometric analysis.

## AUTHOR CONTRIBUTIONS

Conceptualization: H.H.; Methodology: R.H, J.P., Y.L., M-G.A., J.T., C.Z., A.A, D.S., G.R., Y.L., N.H., J.S., L.S., P-Y.S., T.W., J.S., D.W., S.W., K.P., H.H.; Investigation: R.H, J.P., Y.L., J.T., C.Z., A.A, D.S., G.R., N.H., T.W., J.S., S.W., K.P., H.H; Data Curation: R.H, J.P., J.T., C.Z., A.A, D.S., G.R., N.H., K.P., H.H; Supervision: P-Y.S, T.W., J.S., D.W., S.W., K.P., H.H; Manuscript preparation: H.H. wrote the manuscript with review and editing from all authors.

## SUPPLEMENTARY FILE

**Figure S1.**
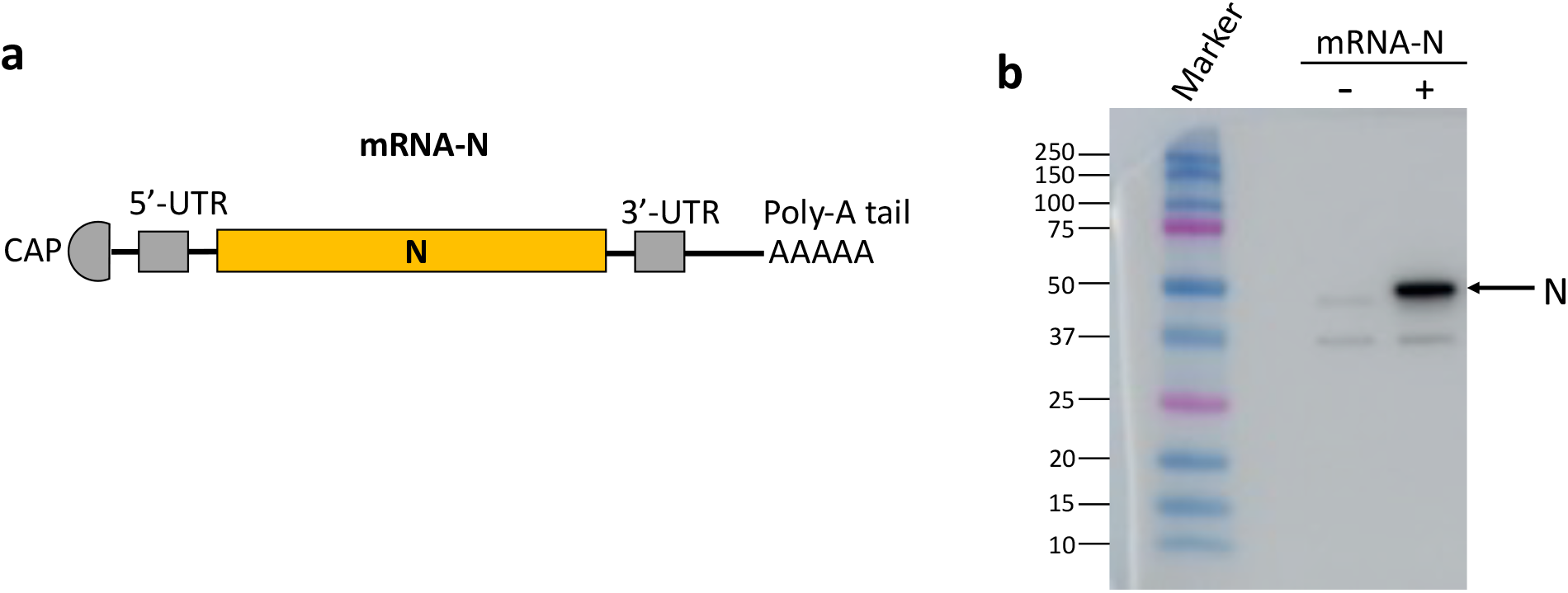
mRNA-N vaccine design and characterization. **(a)** Structure of mRNA-N vaccine. Pseudouridine modified RNA encoding full-length SARS-CoV-2 N protein was synthesized, followed by 5’ capping and 3’ poly-A tailing. (**b)** Western blot confirmation of SARS-CoV-2 N protein expression by mRNA-N. 293T cells were transfected with 2µg mRNA-N-LNP or PBS for 18 hours. Total protein was extracted from the cells for WB analysis. SARS-CoV-2 N protein was detected using a specific anti-N antibody (MA5-29981).

**Figure S2.**
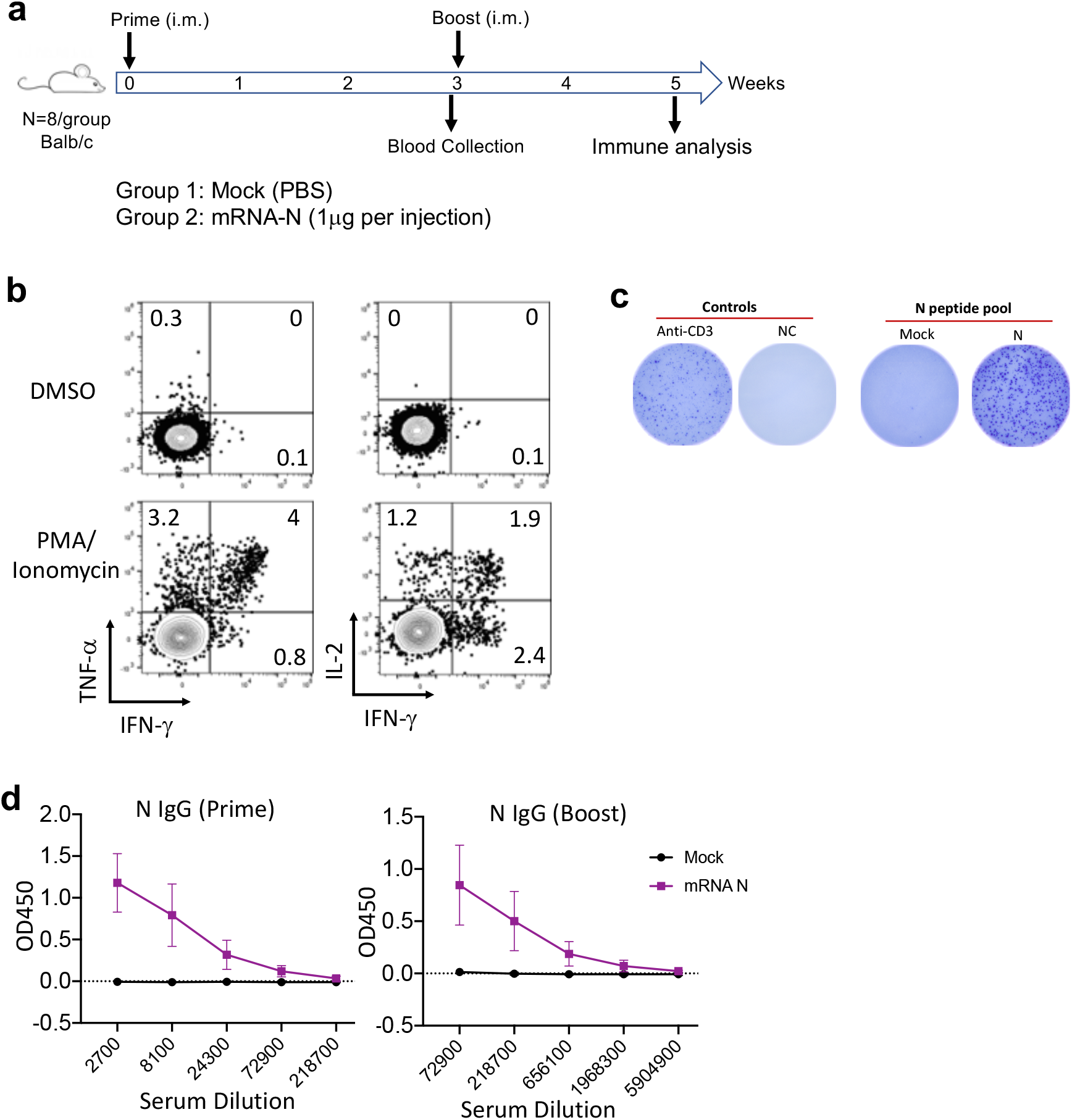
mRNA-N vaccine immunogenicity in mice. **(a)** Vaccination timeline and group design for analysis of mRNA-N vaccine immunogenicity in mice. Balb/c mice (8/group) were intramuscularly (i.m.) vaccinated with mock and mRNA-N vaccine via intramuscular route at week 0 (prime) and week 3 (booster). Blood/serum samples were collected 3 weeks after prime vaccination (before booster) for analysis of vaccine-induced Ab response. 2 weeks after booster vaccination (week 5), all mice were subjected to immune analysis. **(b)** Representative FACS plots for cytokine expression in T cells following DMSO (negative control) or PMA/Ionomycin (positive control) stimulation. **(c)** Representative IFN-γ ELISPOT plots for measurement of N-specific T cells in mouse spleen. Positive control (anti-CD3 stimulation) and negative control (medium only) for the ELISPOT are shown. **(d)** ELISA measurement of N-specific binding IgG in serially diluted (1:3) serum samples. Mean OD values (mean±SD) for serum samples at indicated dilutions after prime (left) and booster (right) vaccination were shown for determination of IgG endpoint titers (EPTs).

**Figure S3.**
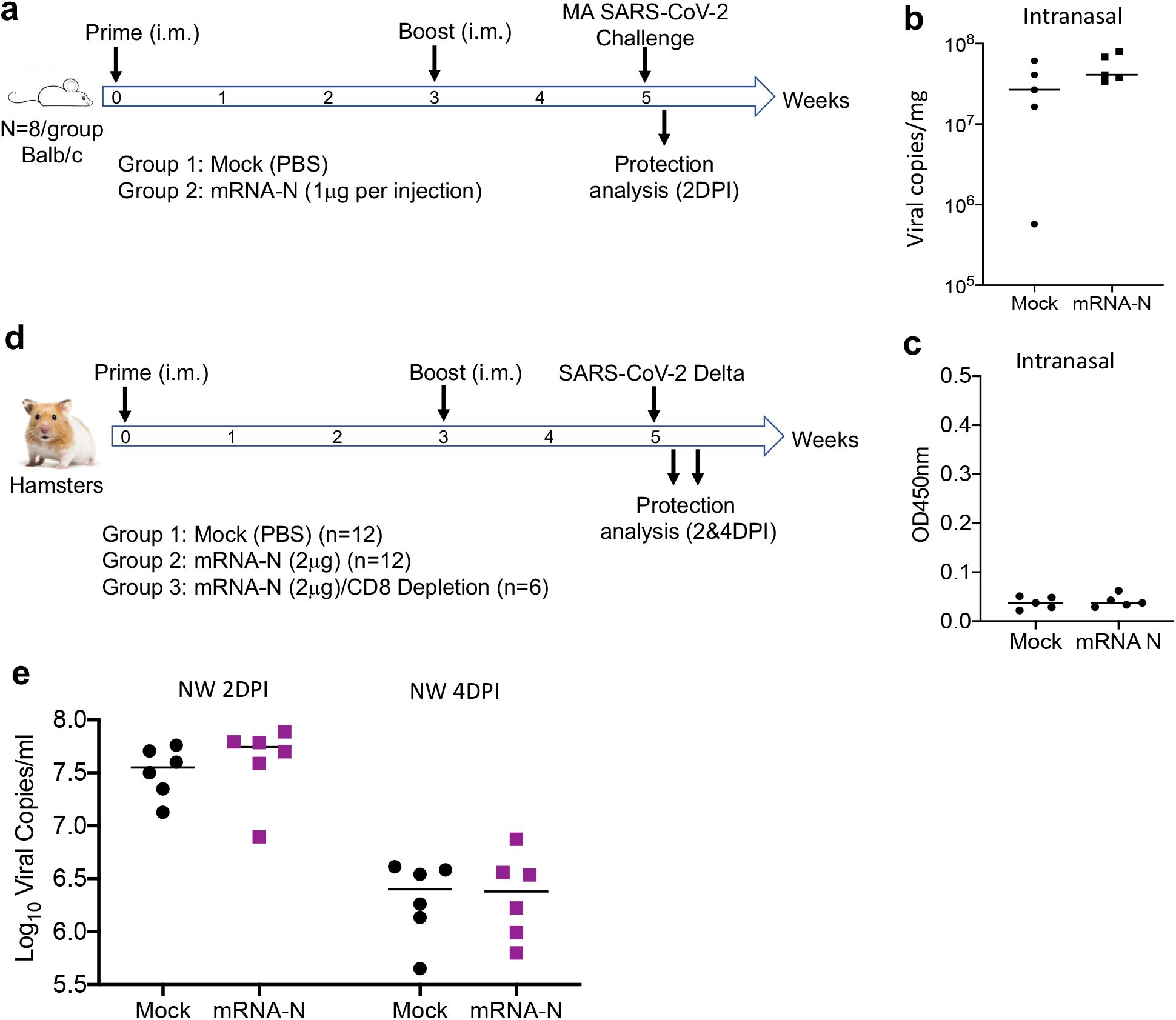
Analysis of mRNA-N-induced protection in mice and hamsters. **(a)** Experimental timeline and group design for analysis of mRNA-N-induced protection in mice following intramuscular (i.m.) vaccination (for Fig. 3a). Two groups of Balb/c mice (8/group) were i.m. vaccinated with mock and mRNA-N vaccine at week 0 and week 3, followed by intranasal challenge with a mouse-adapted SARS-CoV-2 strain (2×10^4^ pfu). 2 days post-infection (2 DPI), viral copies in the lung were analyzed by qPCR. **(b)** Protection analysis in mice following intranasal (i.n.) vaccination. Two groups of Balb/c mice (n=5/group) were i.n. vaccinated with mock and mRNA-N vaccine at week 0 and week 3, followed by intranasal challenge with a mouse-adapted SARS-CoV-2 strain (2×10^4^ pfu). 2 days post-infection (2 DPI), viral copies (Log_10_) in the lung were analyzed by qPCR and were compared between mock and vaccine group. **(c)** Serum antibody response in mice following intranasal (i.n.) vaccination. Sera were collected from the above mentioned, i.n. vaccinated mice 2 weeks after booster vaccination (prior to viral challenge). N-specific binding IgG in sera was measured by ELISA. OD values for individual serum samples (1:30 dilution) were shown. **(d)** Experimental timeline and group design for analysis of mRNA-N-induced protection in hamsters (for Fig. 3b-d). Two groups of hamsters (12/group) were intramuscularly (i.m.) vaccinated with mock (Group 1) or mRNA-N (group 2) at week 0 and week 3, followed by intranasal challenge with the delta strain (2×10^4^ pfu). On 2 DPI (n=6) and 4 DPI (n=6), viral copies in the lung as well as in the nasal washes were analyzed by qPCR. A third group of hamsters (Group 3) (n=6) also received the same mRNA-N vaccine but underwent CD8 depletion three days prior to delta strain challenge. **(e)** Comparison of viral RNA copies (Log_10_) in the nasal washes of hamsters (Group 1 and Group 2) on 2 DPI and 4 DPI in hamsters.

**Figure S4.**
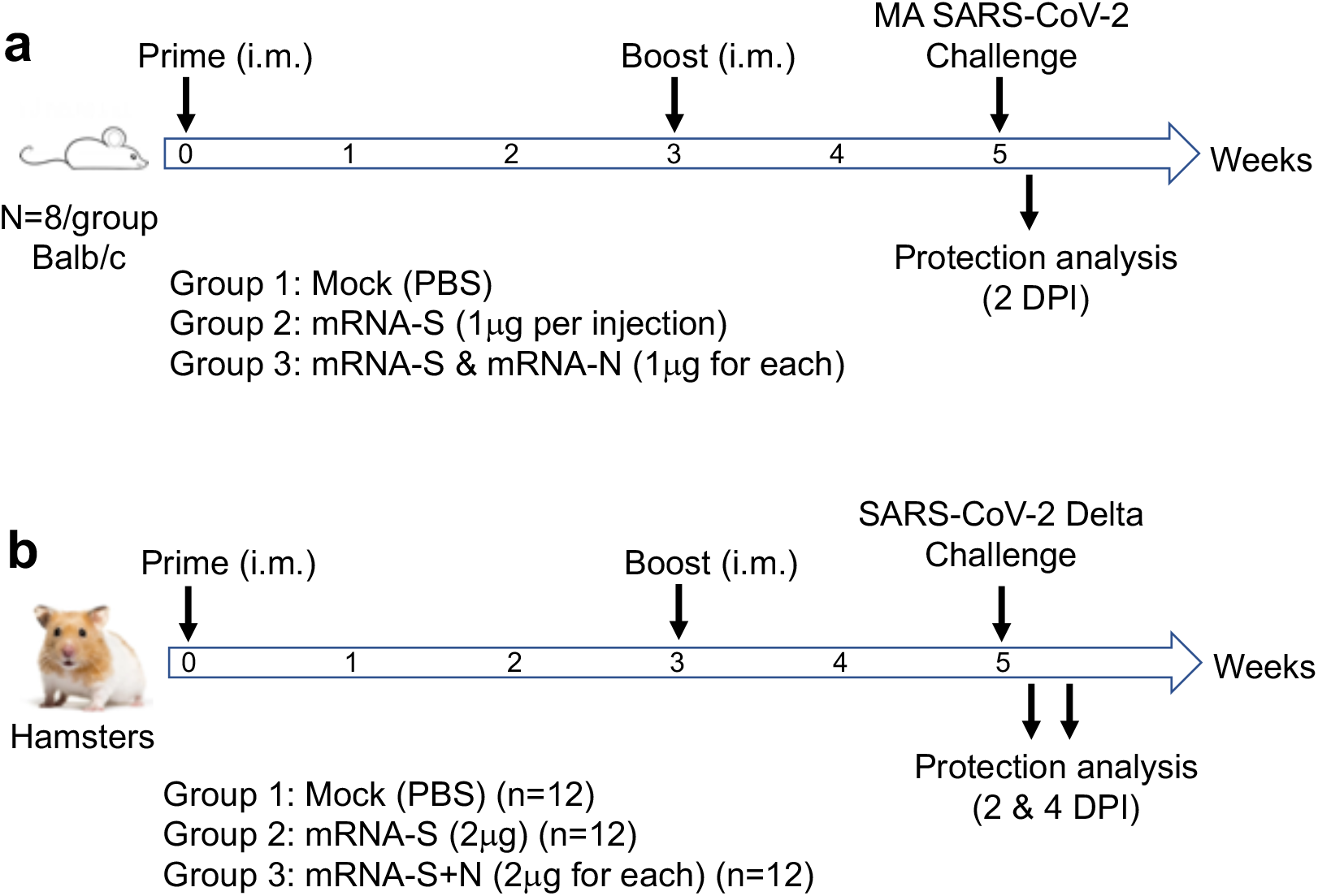
Analysis of mRNA-S and mRNA-S+N induced protection in mice and hamsters. **(a)** Experimental timeline and group design for analysis of vaccine-induced protection in mice. Three groups of Balb/c mice (8/group) were intramuscularly (i.m.) vaccinated with mock, mRNA-S, or combined mRNA-S+N at week 0 and week 3, followed by intranasal challenge with the MA SARS-CoV-2 strain (2×10^4^ pfu). On 2 DPI, viral copies in the lung were analyzed to evaluate vaccine-induced protection. **(b)** Experimental timeline and group design for analysis of vaccine-induced protection in hamsters. Three groups of hamsters (12/group) were intramuscularly (i.m.) vaccinated with mock, mRNA-S, or combined mRNA-S+N at week 0 and week 3, followed by intranasal challenge with the SARS-CoV-2 delta strain (2×10^4^ pfu). On 2 DPI (n=6) and 4 DPI (n=6), viral copies in the lung as well as in the nasal washes were analyzed by qPCR.

**Figure S5.**
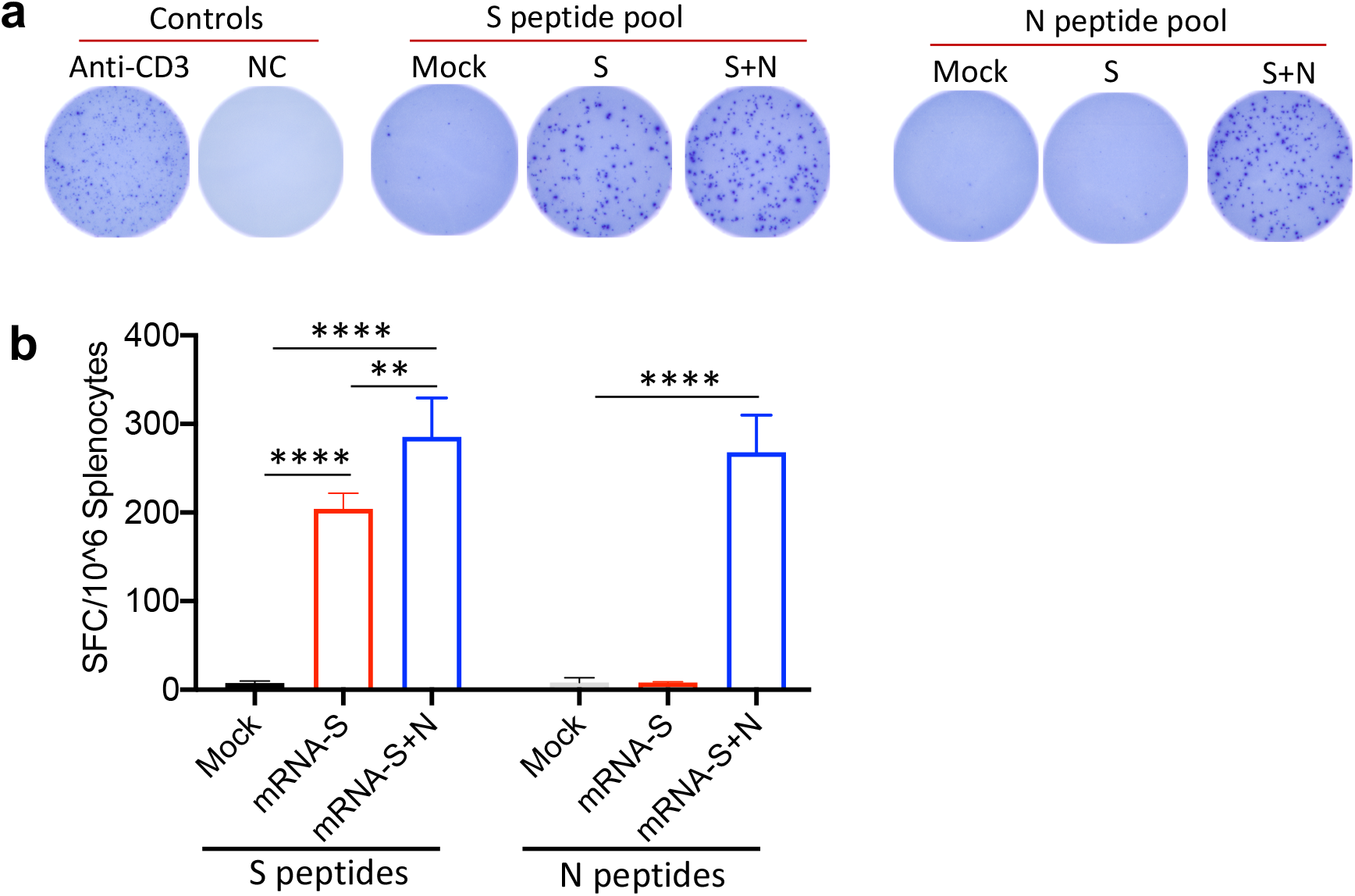
Immune analysis of mRNA-S and combinatorial mRNA-S+N vaccination in mice. **(a)** Representative T-cell ELISPOT plots for measurement of S-specific and N-specific T cells in the mouse spleen following mock, mRNA-S, or combined mRNA-S+N vaccination. Positive control (anti-CD3 stimulation) and negative control (medium only) for the ELISPOT were shown. **(b)** Comparison of S-specific and N-specific T cell response in the mouse spleen following different vaccination based on SFC per 10^6^ splenocytes as determined by ELISPOT. One-way ANOVA was used for statistical analysis. * p<0.05, ** p<0.01, *** p<0.001, **** p<0.0001.

